# Comparative Virome Profiling and Discovery of Novel Viruses in Managed and Wild Bees using RNA-seq

**DOI:** 10.1101/2025.04.27.650873

**Authors:** Zixiao Zhao, M. Catherine Farrell, Rameshwor Pudasaini, Jayla Marvin, Sandra M. Rehan, Hongmei Li-Byarlay

## Abstract

Insect pollinators face growing challenges from viral pathogens that may contribute to population declines, yet comparative data on virome composition in managed versus wild bees remain limited. Using RNA-Seq-based metatranscriptomic analysis, we compared the virome of *Apis mellifera* L. and *Ceratina calcarata* Robertson collected from agricultural and non-agricultural landscapes. We identified multiple known bee pathogens including Black Queen Cell Virus and Deformed Wing Virus in *A. mellifera*, and discovered three novel insect viruses—two iflaviruses and one entomopoxvirus—associated with *C. calcarata*. Notably, one novel iflavirus was identified in *A. mellifera*. Our findings reveal species- and landscape-specific viral prevalence, highlight viral diversity beyond canonical bee pathogens, and provide genomic resources for future functional studies. These data offer new insights into virus-host dynamics and underscore the utility of virome profiling in understanding pollinator health at the molecular level.

## Introduction

Animal pollinators have critical ecological functions in maintaining the diversity of global ecosystems [1, 2]. About 87.5% of flowering plants rely on animal pollinators for successful reproduction [3]. In agriculture, approximately 75% of crop species require pollinators, contributing about 30-35% of total crop production [4, 5]. Bees (Hymenoptera: Apidae) are considered the most abundant and important pollinators globally. Honey bee species (*Apis* spp.), with a global distribution due to domestication, contribute about 80% of total pollination and create about $700 million value annually in the US [6]. However, the ecological significance of non-*Apis* wild bee species shouldn’t be neglected. Wild bees provide pollination services to many crops, even when managed honey bees are not available [7, 8]. Among the over 17,000 wild bee species worldwide, with a wide range of social structures from eusocial to solitary, eusocial species may be more susceptible to viral infections due to their high colony densities, frequent social interactions and genetic factors [19]. For last several decades, both wild and managed honey bees are suffering from colony declines [9-12], which not only causes economical losses to apiary industry, but more importantly, reduces pollination service and imposes potential risks for global food security.

The causes of bee populations decline are complex. Factors responsible for bee populations decline include habitat loss and land fragmentation, increased use of chemical pesticides and other agrochemicals, introduction of pests and pathogens, and climate change [11, 13]. Among these stressors, pathogenic viruses are considered the direct and one of the most significant hazards to both managed and wild bee populations (Ullah et al. 2020). Currently identified bee pathogenic viruses belong to positive-sense single-stranded RNA (+ss RNA) virus category. The majority of the pathogenic bee viruses are from two families under the order Picornavirales: Iflaviridae and Dicistroviridae. Iflaviridae has only one genus: *Iflavirus*, which includes Sacbrood virus (SBV), Deformed wing virus (DWV), Varroa destructor virus-1 (VDV-1), and Slow bee paralysis virus (SBPV). While the family Dicistroviridae includes Black queen cell virus (BQCV), Kashmir bee virus (KBV), Acute bee paralysis virus (ABPV), and Israeli acute paralysis virus (IAPV) [14]. Two additional pathogenic viruses of bees are Chronic bee paralysis virus (CBPV), an unclassified virus with two genomic RNA segments [15], and Lake Sinai viruses (LSVs), members of the order Nodamuvirales [14], with eight identified lineages [16]. The symptoms caused by pathogenic viruses vary, including body deformities, color changes, uncapped cells with larvae or pupae, paralysis of adults, which eventually increase mortality and lead to colony degradation. IAPV infections have been discovered association with colony collapse disorder (CCD), which responsible for large-scale honey bee colony loss in the US [17].

Despite managed honey bees, the pathogenic viruses can also infect wild bee species [18-20]. For example, DWV has been reported to infect a wide range of hymenopterans, including some social and solitary bees as well as several wasp species [18, 20, 21]. Dolezal et al. [19] reported DWV and SBV were widely detected in wild bees collected in Iowa, while IAPV, LSV and BQCV were also detected in wild be species. Furthermore, DWV, BQCV and SBV can be deposited to pollen and remain infective, which can contribute to inter-taxa transmission among bees, and wasps [20]. These studies suggested viruses can be a potential factor responsible for colony declines in wild bee species.

Although agriculture provides important foraging resources for bees, it is also considered one of the major reasons for colony declines. It has converted natural ecosystems to farmland primary for food and fiber productions. This process is accompanied by loss of biodiversity and use of agrochemicals causing habitat loss, fragmentation and degradation. Agriculture can also alter host-pathogen interactions at multiple levels. A study demonstrated that the relevance of plant viruses is negatively correlated with the biodiversity in agricultural landscape [22]. However, whether agricultural activities affect viral prevalence in insects, especially in honey bees and other bee species, is still unknown.

The small carpenter bee, *Ceratina calcarata* Robertson (Hymenoptera: Apidae), is a wild bee species native to the eastern part of North America. It is considered as an indicator species of healthy ecosystems and important pollinators for several native and agricultural plants [23]. Due to the feature of subsociality, *C. calcarata* is used for studying pollinator ecology, behavior, evolution, genomics and nutrition [24-27]. However, little is known about the virology of this species. The impact of viruses on *C. calcarata* populations and whether the viral prevalence is related to agricultural activities are not clear.

Here we presented our metagenomic studies on viruses associated with western honey bee (*A. mellifera* L.) and *C. calcarata* in response to different agricultural intensifications. Bees were collected from three landscape types representing levels of agricultural intensification: conventional farming, organic farming and roadside with no agricultural activities. The viral RNA was extracted from collected bee samples and subjected to high-throughput sequencing. We analyzed known and novel viruses associated with both bee species, which were highly specific to location and bee species. Our study provides a new methodology and unique knowledge for bee-virus interactions.

## Results

### Reconstruct virome from viral RNA-Seq reads

The RNA-Seq of extracted viral RNA generates approximately 17 to 36 million raw read pairs for each *A. mellifera* and *C. calcarata* sample. After quality trimming and partially removing reads from respective host genome (Amel HAv3.1 PRJNA471592 and Ccalc.v3 PRJNA791561), 5 to 25 million pairs were kept for each sample (**Supplementary file S1**). Trinity *de nove* assembled 366,289 contigs for *A. mellifera* and 203,317 contigs for *C. calcarata* virome, which consists of 188,433 and 91,314 non-redundant clusters. The assembled contigs were subject to BLASTN searches against non-redundant databases (nr), which claimed 541 clusters and 1187 contigs from *A. mellifera* assembly potentially belong to viruses. In contrast, 62 clusters and 84 contigs from *C. calcarata* assembly were potentially from viruses. A large portion of assembled contigs were from either host genes or the plants they have visited.

### Viruses infecting bees and other insects detected in A. mellifera and C. calcarata virome assemblies

BLASTN search identified four known bee pathogenic viruses from *A. mellifera* virome: BQCV, DWV, SBV, and LSV3 with fully assembled genome sequences. Two other members of the LSV group: LSV1 and LSV2 were detected. But their sequences were only partially assembled. Positive and negative strands were detected in all of the above viruses except LSV2, from which only positive strand was detected. We also detected *Apis mellifera* filamentous virus (AmFV) [28] and Hubei partiti-like virus 34 [29] in *A. mellifera*. The AmFV were partially assembled and the Hubei partiti-like virus 34 demonstrated a fully assembled genome (**Table 1A**). In *C. calcarata*, BQCV was the only detected bee pathogenic virus and appeared as partial-assembled as negative-strand contig (**Table 1B**).

**Table 1.**
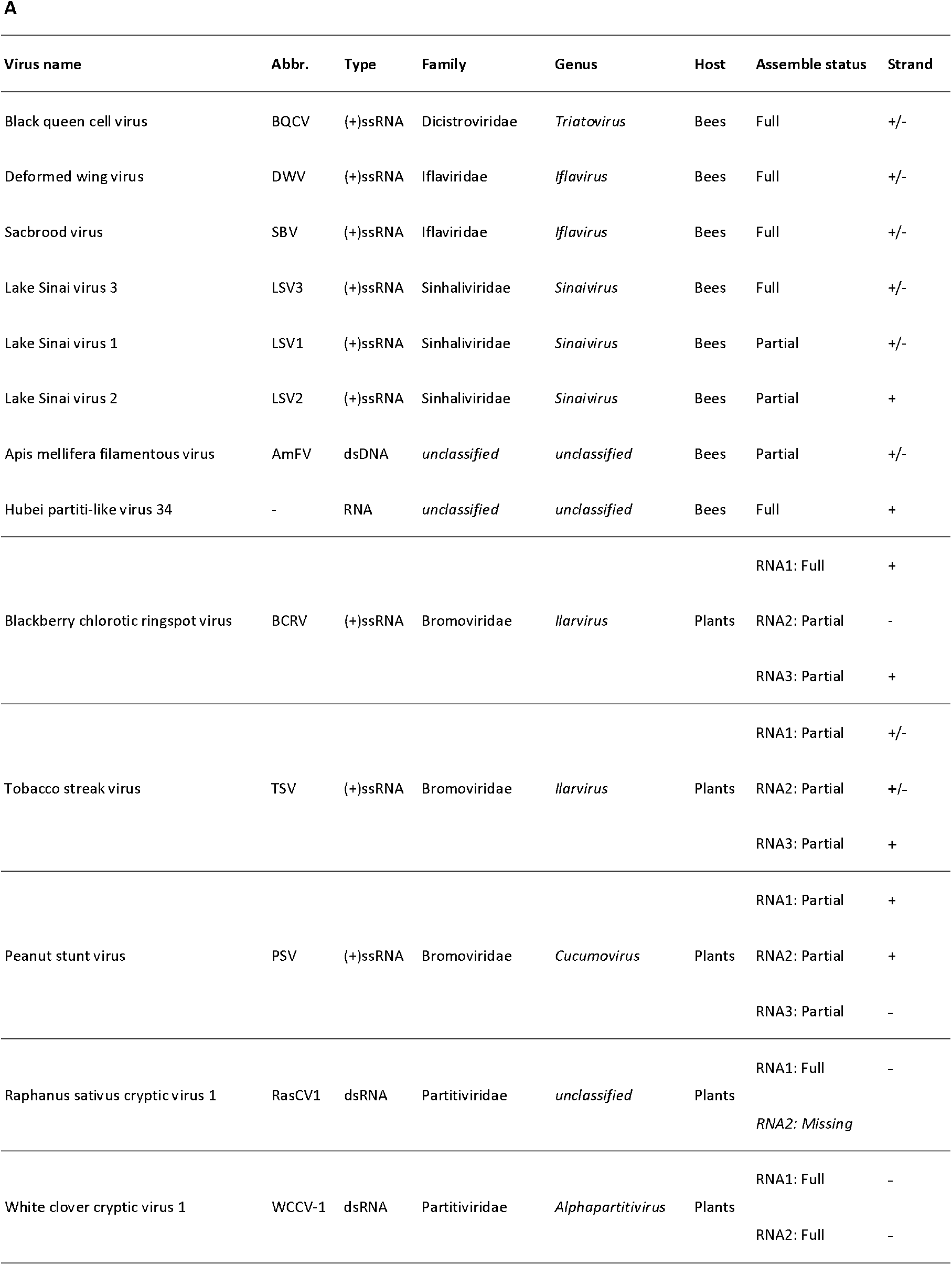
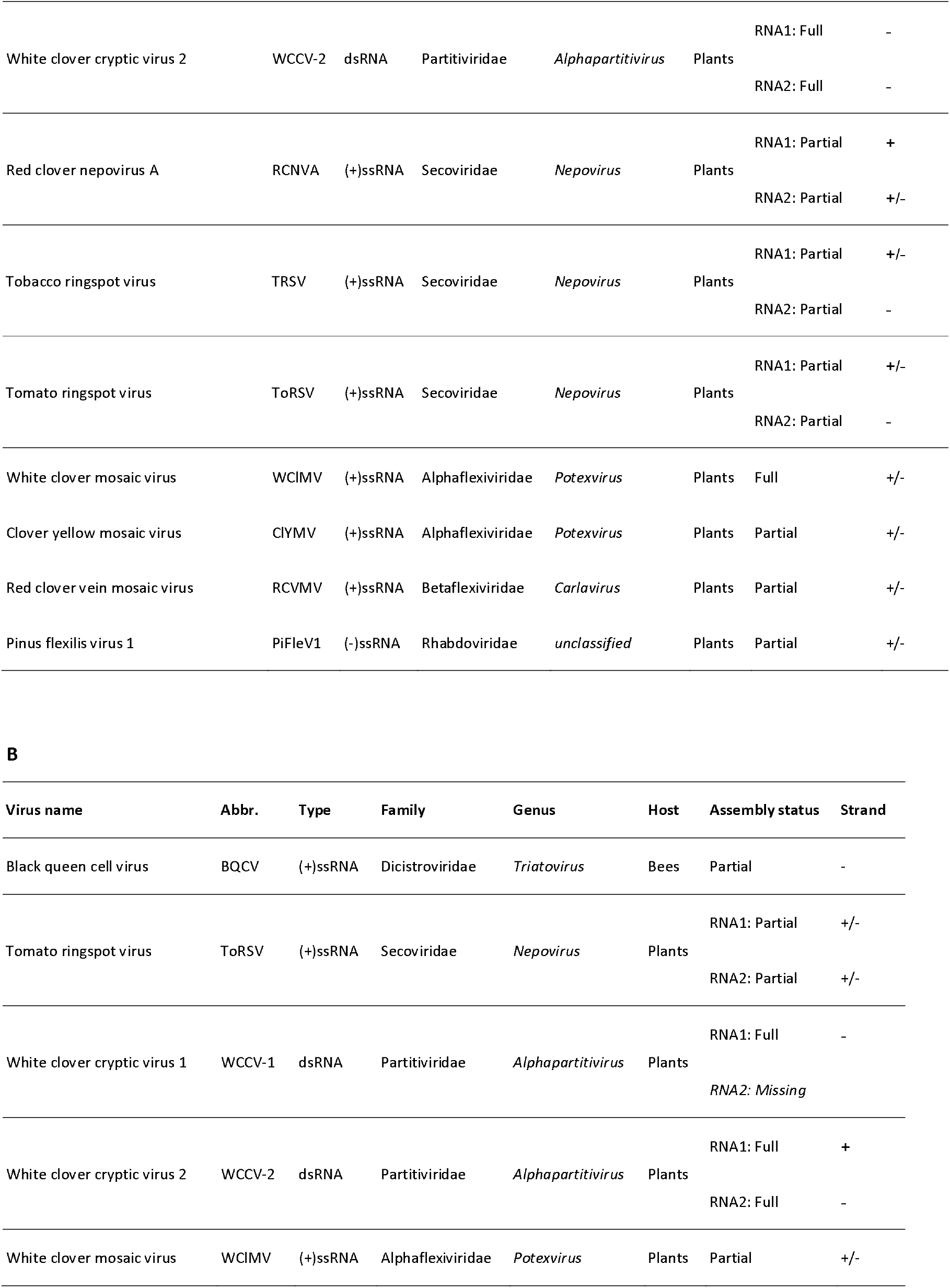
List of identified insect and plant viruses in **(A)** *A. mellifera* and **(B)** *C. calcarata* virome assemblies, assembled from RNA-Seq data.

**Table 2.**
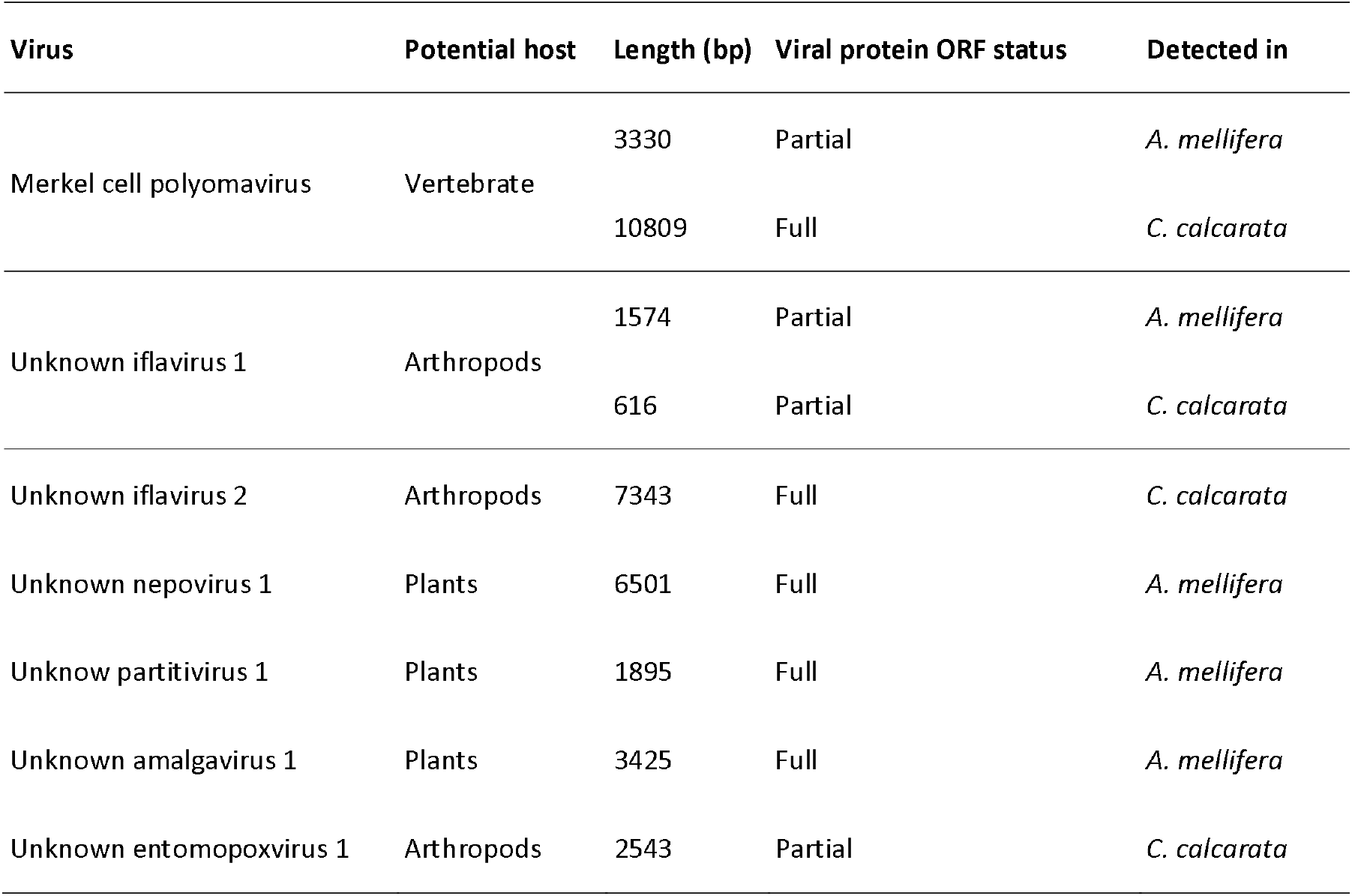
List of novel viruses detected in *A. mellifera and C. calcarata* virome assemblies assembled from RNA-Seq data.

### Plant viruses discovered in A. mellifera and C. calcarata virome assemblies

A range of plant viruses were detected in virome assemblies of both bee species. In *A. mellifera*, white clover cryptic virus 1 (WCCV-1) and white clover cryptic virus 2 (WCCV-2) from genus *Alphapartitivirus* (family: Partitiviridae, +ssRNA), as well as white clover mosaic virus (WCMV) from genus *Potexvirus* (family: Alphaflexiviridae, +ssRNA) were fully assembled. Blackberry chlorotic ringspot virus (BCRV) from genus *Ilarvirus* (family: Bromoviridae, +ssRNA) and *Raphanus sativus* cryptic virus 1 (RsPV1) from Partitiviridae (+ssRNA) have one genomic segment fully assembled, while other segments were either partially assembled or completely missing. Partially assembled plant viruses include 7 +ssRNA viruses: peanut stunt virus (PSV) and tobacco ringspot virus (TRSV) from Bromoviridae, clover yellow mosaic virus (ClYMV) from Alphaflexiviridae, red clover nepovirus A (RCNVA), tobacco streak virus (TSV), and tomato ringspot virus (ToRSV) from genus *Nepovirus* (family: Secoviridae), red clover vein mosaic virus (RCVMV) from Betaflexiviridae, and one -ssRNA virus: *Pinus flexilis* virus 1 (PiFleV1) from Rhabdoviridae (**Table 1A**). However, the plant viruses in *C. calcarata* virome were considerably less diverse compared to *A. mellifera*. WCCV-2 was the only fully assembled plant virus, while WCCV-1, BCRV, ToRSV, and WClMV were partially assembled in *C. calcarata* (**Table 1B**).

### Novel viruses in A. mellifera and C. calcarata virome assemblies

The genus *Iflavirus* (family: Iflaviridae) consists of +ssRNA viruses infecting arthropods, including three bee pathogenic viruses SBV, DWV, and SBPV. Two novel viruses under genus *Iflavirus*, referred to as unknown iflavirus 1 and unknown iflavirus 2, were identified in *C. calcarata* virome assembly as positive strands. The phylogenetic analysis using translated polyprotein amino acid sequence suggests both viruses are in the same group as SBV. The unknown iflavirus-1 has the highest identity with Darwin bee virus-2, an unclassified picornavirus infecting honey bee [30] (**Figure 1A**). This virus was also detected in *A. mellifera* virome assembly as a negative strand, suggesting that it can infect and replicate in honey bees.

**Figure 1.**
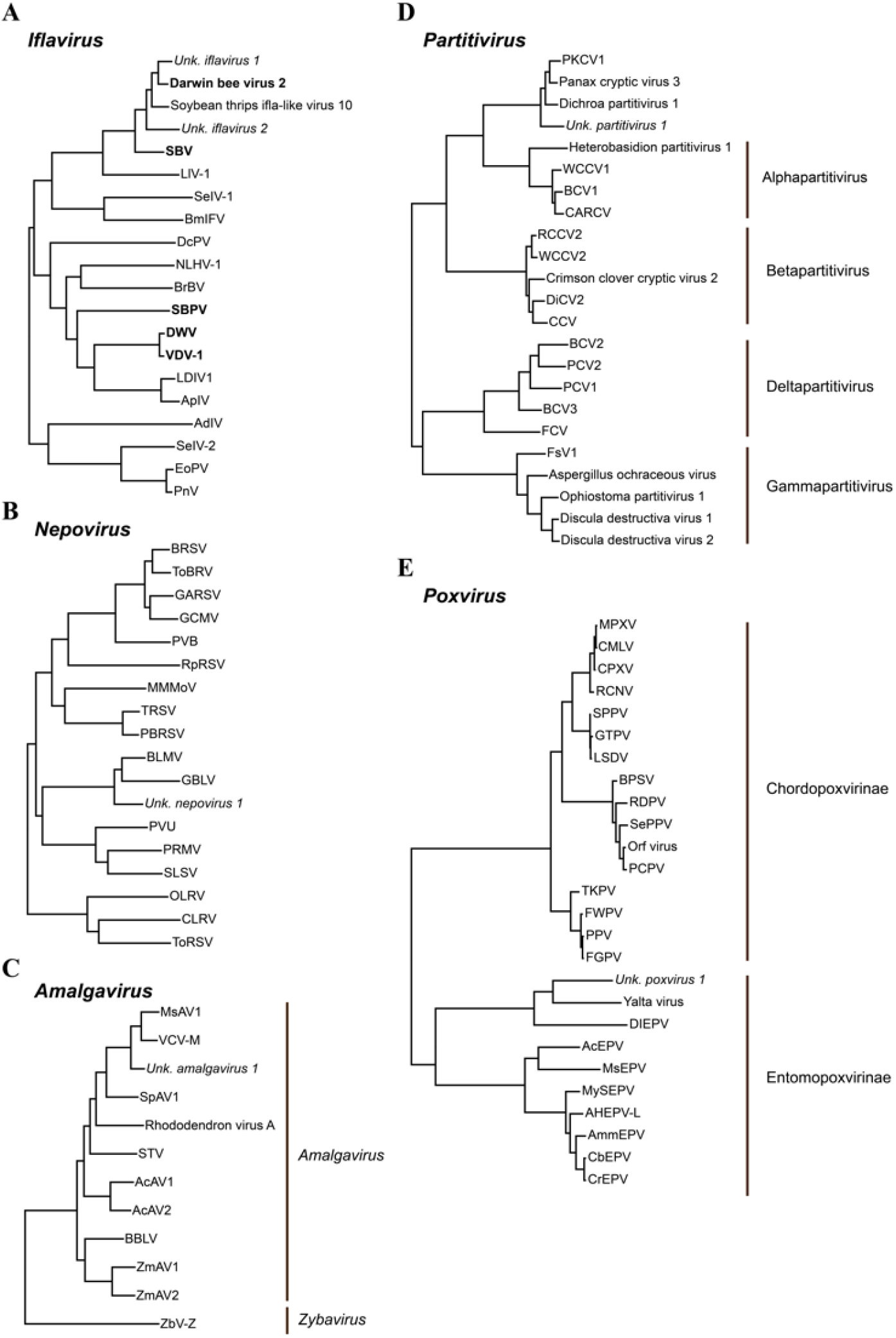
Phylogenetic trees of novel viruses discovered from *A. mellifera* and *C. calcarata* virome assemblies by Maximum-likelihood analysis of sequences of viral proteins: **(A)** polyprotein (iflavirus), **(B)** polyprotein from RNA2 (neprovirus), **(C)** RNA-dependent RNA polymerase (amalgavirus), **(D**) RNA-dependent RNA polymerase from RNA1 (partitivirus), and **(E)** RNA-dependent RNA polymerase submit RPO132 (poxvirus). The names in bold correspond to honeybee-associated viruses. Abbreviations and protein accessions of known viruses are included in **Supplementary File S2**.

In *C. calcarata*, we detected a novel poxvirus (a double-strand DNA virus). Its DNA-dependent RNA polymerase RPO132 is partially assembled. Phylogenetic analysis showed this virus is closely related to the Yalta virus, an uncategorized poxvirus from *Drosophila melanogaster* and *Diachasmimorpha* entomopoxvirus (DIEPV), from parasitoid wasp *Diachasmimorpha longicaudata*. The three viruses form a sister cluster to other entomopoxviruses (EPVs) (genus: *Alphaentomopoxvirus, Betaentomopoxvirus*, and *Deltaentomopoxvirus*), while all the vertebrate poxviruses (genus: *Avipoxvirus, Capripoxvirus*, and *Parapoxvirus*) are clustered into a separate group. The result suggests this novel virus belongs to the subfamily Entomopoxvirinae (EPV) that primarily infect insects (**Figure 1E**).

Three additional novel plant viruses were detected in the *A. mellifera* virome, including one from the genus *Nepovirus* (unknown nepovirus 1, family: Secoviridae), one from family Partitiviridae (unknown parititivirus 1, family: Partitiviridae), and one of genus *Amalgavirus* (unknown Amalgavirus 1, family: Amalgaviridae). Nepovirus has two positive-strand genomic RNAs [31]. For unknown nepovirus 1, the open reading frame (ORF) of polyprotein RNA2 was fully assembled. However, the correspondent RNA1 encoding viral coat protein was not detected. The phylogeny of the polyprotein sequence shows that it is most closely related to Blueberry leaf mottle virus and Grapevine Bulgarian latent virus, forming a separate cluster (**Figure 1B**). Similarly, only RNA1 encoding RNA-dependent RNA polymerase (RdRp) was fully assembled for another potential plant virus: unknown parititivirus 1. The phylogenetic analysis of RdRp showed that it is clustered with several uncategorized partitivirus viruses associated with plant: *Polygonatum kingianum* cryptic virus 1, Panax cryptic virus 3, and *Dichroa partitivirus* 1. This cluster is a sister group to plant-infecting genus *Alphapartitivirus* (**Figure 1D**). The genome of unknown amalgavirus 1 was fully assembled with two complete ORF: the ORF1+2 fusion protein responsible for genome replication, and a coat protein. The phylogeny of fusion proteins showed that it is the most closed to Vicia cryptic virus M (VCV-M) and Medicago sativa amalgavirus 1 (MsAV1) and have a clear separation with *Zygosaccharomyces bailii* virus Z (ZbV-Z), a yeast-infecting virus belongs to the genus *Zybavirus* (**Figure 1C**).

In addition, other viruses, including phages and a human virus: Merkel cell polyomavirus (Family: Polyomaviridae) were identified in both virome assemblies. The viral genome consists of circular double-stranded RNA (dsRNA), which resulted in the assembly of two identical full-length viral sequences into one contig in *C. calcarata* virome. In *A. mellifera*, only partial sequence was assembled. How the Merkel cell polyomavirus was introduced into virome and its effects in both species, are still unknown.

### Abundance of viruses in response to different agricultural landscapes

The distributions of the above detected viruses were highly site-specific. In *A. mellifera*, among all identified insect viruses, five known bee viruses: BQCV, DWV, LSVs, SBV, and AmFV were detected at all of the experimental sites regardless of agricultural practices, with AmFV only marginally detected in conventional farm C1. In contrast, the novel unknown ilfavirus 1 was only detected in samples from organic farm O2 (**Figure 2A**). For five known bee viruses, the edgeR did not show any significant differences at virus level. However, a separate 2834-bp LSV3 segment, containing ORFs of two viral proteins, was significantly increased in bees collected from organic sites compared to bees collected from roadside site (FDR: 0.04). The Hubei partiti-like virus was also identified at all sites except conventional farm C1 (**Figure 2A**). Its full-length sequence was significantly higher in bee samples collected from organic farms compared to bee samples collected from both conventional and roadside sites (FDR: 0.006 to conventional, 0.003 to roadside).

**Figure 2.**
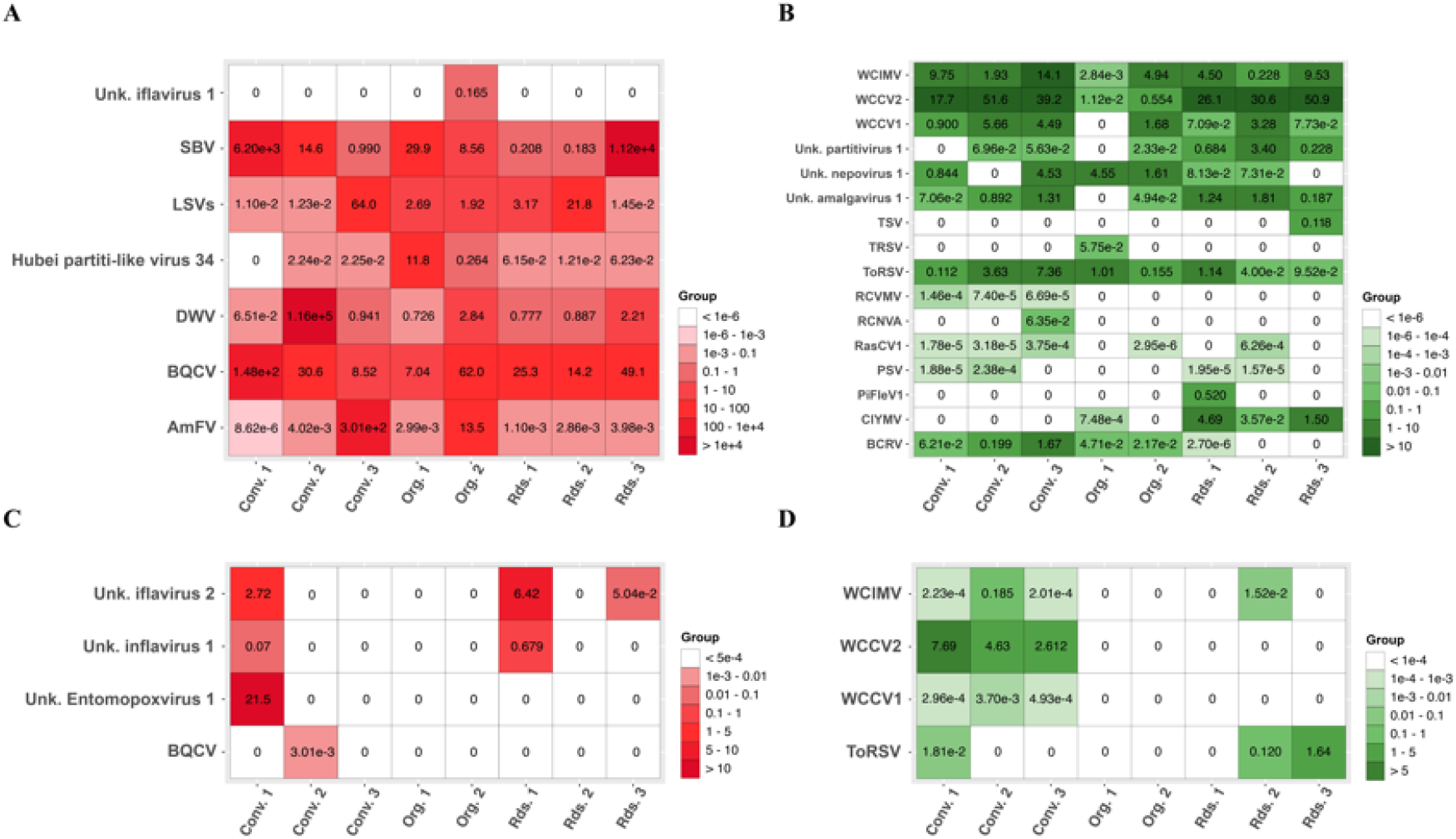
Heatmaps demonstrating the relative abundance of **(A)** insect viruses in *A. mellifera*, **(B)** plant viruses in *A. mellifera*, **(C)** insect viruses in C. *calcarata*, and **(D)** plant viruses in *C. calcarata* at each sampling site. Average normalized count-per-million reads (cpm) from RNA-Seq was calculated for each virus of each site and used in the heatmap. White color indicates the virus is undetected on that site. Potential insect and plant viruses were included.

The presence and levels of plant viruses were highly dependent on the sampling locations. For instance, TSV and PiFleV1 were detected only at one separate roadside site each, while RCNVA and TRSV were found only at one conventional and one organic site, respectively. RCVMV was identified across three conventional sites, but at relatively low levels. WCIMV, WCCV-2, and ToRSV were expressed at all experimental sites (**Figure 2B**). WCCV-1 was the only significant plant virus detected by edgeR. The levels of both genome segments decreased significantly in organic sites, compared to both conventional (FDR, RNA1: 0.011, RNA2: 0.006) and roadside sites (FDR, RNA1: 0.006, RNA2: 0.003). The viral levels of BRCV at roadside sites were either very low or undetected, but the overall level is too low to perform statistical analysis in edgeR.

*C. calcarata* hosted relatively fewer viruses compared to *A. mellifera*. The presence of insect viruses was also highly site-specific. BQCV and unknown entomopoxvirus 1 were only found at one conventional site, whereas two novel ilfavirus were detected in roadside sites and conventional farm C1. None of the viruses were detected from bee samples collected from organic sites, conventional farm C3, and roadside R2 (**Figure 2C**). Regarding the plant viruses, WCCV-1 and WCCV-2 had a similar pattern, being detected only at conventional farms, while WCIMV and ToRSV were detected in few bee samples collected from roadside sites as well (**Figure 2D**). However, their levels were not statistically significant. Similarly, none of the plant viruses were detected in bee samples collected from organic sites.

## Discussion

The metagenomic approach based on viral RNA-Seq was adopted for identifying and quantifying known viruses, as well as discovering novel viruses. The present study surveyed the abundance of viruses associated with two bee species. Findings show that the virome compositions were site-specific. However, the *A. mellifera* carried a wider range of both insect and plant viruses than *C. calcarata*. The results also indicate that the *A. mellifera* may transmit plant diseases more effectively than *C. calcarata*.

### The Honey bee viruses in A. mellifera and spillover *to C. calcarata*

The levels of BQCV, DWV and SBV were in high levels in *A. mellifera* virome at all experimental sites, suggesting the high prevalence of these viruses in *A. mellifera*. In contrast, none of the acute bee paralysis virus complex (ABPV, KBV and IAPV) were detected. This result is consistent with other studies in the Northeast of the US [20, 32]. LSV is a highly variable, multi-strain bee virus that does not have any significant symptoms but is likely to weaken colonies. The LSV contigs assembled in this study are most closely related to LSV1, LSV2 and LSV3. The LSV3 was detected from all experimental sites and the only LSV with full-length genome sequence assembled. The LSV3 was originally identified in Slovenia [33] and has been detected in the Western US [34]. The study confirmed LSV3 has been spread to the New England area, in the US.

Two known viruses associated with *A. mellifera* were also detected: AmFV and Hubei partiti-like virus 34. The AmFV is a weak pathogen in *A. mellifera* and rarely causes significant symptoms. Its infection has been reported related to BQCV and DWV [28, 35]. We detected AmFV from all sites, while its viral loads were from extremely low at conventional farm C1 to medium-high at conventional farm C3. The Hubei partiti-like virus 34 was first discovered as an arthropod RNA virus [36] and was reported associated with hymenopteran species, including *A. mellifera* and Asian hornet (*Vespa velutina*) in East Asia and South Europe [29, 37]. The current study reported the low to moderate infection of Hubei partiti-like virus 34 from all sites except conventional farm C3 site. The potential impact of this virus on honey bee health is still unknown.

The spillover of managed honey bee pathogen to wild bees has been considered as a threat to native bee populations [38, 39]. In genus *Ceratina*, DWV was found spillover to two species *C. dupla* [20] and *C. smaragdula* [40]. The current study showed that, given the high prevalence of both BQCV and DWV in *A. mellifera*, BQCV may have spilled over to *C. calcarata*, while no evidence of DWV spillover was observed. BQCV was only detected from conventional farm C2 in relatively low prevalence and viral load, indicating that this spillover to *C. calcarata* is not a common scenario.

### Potential viruses infecting A. mellifera and C. calcarata

The primary hosts for iflaviruses are arthropods. We discovered two closely related novel ifla-like viruses. Unknown iflavirus 1 was detected in both *A. mellifera* and *C. calcarata*, while the unknown iflavirus 2 was detected only in *C. calcarata*. The prevalence of both viruses was low, however, the viral load of unknown ilfavirus 2 was slightly higher in *C. calcarata* samples collected from conventional farm C1 and roadside site 1, compared to the unknown ilfavirus 1 in the same samples.

The EPVs are double-stranded DNA (dsDNA) viruses that infect insects and predominantly insect pathogens. The unknown entomopoxivirus 1 detected in *C. calcarata* are more closely related to Yalta viruses and DIEPV than other entomopathogenic viruses. Although the role of Yalta virus is still unknown, the DIEPV has been found replicated in both parasitoid wasp *D. longicaudata* and its host fruit fly (*Anastrepha suspensa)*. The replication of virus is highly virulent to the host fruit fly and no negative impact to wasp host, suggesting a mutualistic symbiotic relationship of DIEPV to parasitoid wasp [41]. Although both *C. calcarata* and *D. longicaudata* are from order Hymenoptera, no such parasitoid relationship was observed in *C. calcarata*. Whether this novel entomopoxivirus primary infects *C. calcarata* or just an overspill event to *C. calcarata* yet to be studied.

Although our findings reveal the presence of viral sequences in bee samples, it does not confirm active infection or pathogenicity in these bees. Some of the viral sequences may originate from environmental exposure, gut contents, or external body surfaces, rather than systemic infection. Therefore, the present study does not infer a direct relationship between viral presence and bee health or stress levels (Olgun et al., 2020).

### Bees carrying plant viruses

Many plant viruses can travel within or attach to pollen grains [42, 43], including BRCV, TSV, PSV, WCCV-1, WCCV-2, TRSV, ToRSV, and WClMV as observed in the present study However, only TSV, TRSV, and ToRSV have been proven vertically and horizontally transmitted by pollen. The rest were only detected by RNA-Seq [43]. These findings show that pollinators such as *A. mellifera* posing potential risks due to these viruses. However, none of the detected viruses were on the list of known honey bee-transmitted viruses, according to Fetters et al [43]. TRSV was reported infect honey bee [44], however whether cross-kingdom infection assist transmitting diseases is unknown. On the other hand, bees are a powerful tool for monitoring plant diseases [45]. The plant viruses detected in *A. mellifera* resembles to the results of other metagenomic studies of bees and pollen at family and order levels [45, 46], while the species composition is determined by plant and prevalence of plant viruses at the location.

Compared to *A. mellifera*, plant viruses in *C. calcarata* are much less diverse. Previous studies have shown that pollen from clover and rose is a major food resource for bees [47], and white clover pollen significantly increases the fitness of *C. calcarata* [23]. Consistent with that, among four plant viruses detected in *C. calcarata* virome, WCCV-1, WCCV-2, WClMV primarily infect white clover (*Trifolium repens*), and the host range of ToRSV include rose and red clover (*T. pratense*). None of the detected viruses originated beyond *C. calcarata’s* foraging range, suggesting that *C. calcarata* may be an ideal bee species for monitoring white clover-associated pathogens within localized areas.

## Material and method

### Sample collection

Both honey bee (*A. mellifera*) and small carpenter bee (*C. calcarata*) samples were collected from sites representing three different types of land use: conventional farm, organic farm and roadside. Honey bees and the small carpenter bees were collected by sweep nets from May to July 2018 in Strafford County, New Hampshire. We collected both species within three conventional farms and three organic farms. The roadside samples were collected in the urbanized area within the University of New Hampshire campus. The detailed about locations and GPS coordinates are described in previous publications [48, 49]. At least ???? bees were collected at each sampling site, and to minimize age-related variation, all bee samples were collected from forager bees. Bee samples were flash-frozen in liquid nitrogen and temporarily stored at -80 °C.

### RNA extraction, sequencing and data cleaning

The viral RNA was extracted from *A. mellifera* and *C. calcarata* individually by Quick-RNA viral kit (ZYMO RESEARCH, Irvine, CA). DNase I (ZYMO RESEARCH) in-column digestions were applied to remove genomic and environmental DNA. For mRNA library generation, 200ng of total RNA was treated with NEBNext Poly mRNA Magnetic Isolation Module E7490L (New England Biolabs, USA) following the manufacturer protocol. Subsequently, the isolated mRNA was fragmented for 10 minutes. cDNA was synthesized and amplified for 12 PCR cycles using NEBNext Ultra II Directional (stranded) RNA Library Prep Kit E7760L for Illumina (New England Biolabs, USA) with NEBNext Multilex oligos indexes kit E6440S following the manufacturer directions. Distributions of the template length and adapter-dimer contamination were assessed using an Agilent 2100 Bioanalyzer and High Sensitivity DNA kit (Agilent Technologies, Inc). The concentration of cDNA libraries was determined using Invitrogen Qubit dsDNA HS reagents and read on a Qubit Fluorometer (Thermo Fisher), and the cDNA libraries were paired-end 150bp format sequenced on a Novaseq SP system (Illumina, San Diego, CA). In total, 17 honey bee samples and 16 *C. calcarata* samples were sequenced (**Supplementary File S1**). Raw reads were subject to adapter and quality trimming by Trimmomatic [50]. Sample metadata from collection was used to assign each sample to either *A. mellifera* or *C. calcarata*. To remove endogenous transcripts from bee hosts, the reads were respectively aligned to honey bee and small carpenter bee [25] genomes (including mitochondrial sequences) using Bowtie2 [51]. Read pairs that were concordantly aligned to bee genomes were considered as endogenous transcripts and removed from further analysis.

### Virome assembly and viral identification

To reconstruct the virome of *A. mellifera* or *C. calcarata*, we first merged all the pre-cleaned reads from the same organism to increase the sequence coverage. Then two *de novo* assembler: Trinity [52] and metaSPAdes [53] were used with default setting. The results were comparable; however, the total running time for Trinity was lower. Therefore, we chose Trinity for the subsequent analyses.

Despite the pre-cleaning process, the assembled virome still contains host sequences. To identify virus sequences, we used BLASTN to search assembled contigs to default nr/nt database with E-value cutoff of 1e-7. Contigs with first three top hits to viral sequence from database was retained as viral contigs. For further virus identification, we examined the blast results of contigs >500 nt. A contig was assigned as a known virus if it aligned stringently (E-value = 0) to existing viruses. If the alignment is less stringent (0<E-value<1e-7), it was assigned as a novel virus. The “full-length” or “partial” labels were based on the completeness and intactness of viral protein of each viral contig. For potential novel viral contigs, ORF finder and BLASTP were used to translate and predict viral protein. MEGA 11 [54] was used to construct Maximum-likelihood phylogenetic tree based on the sequence of homologous proteins of known and novel viruses.

### Abundance of viruses and differential analysis between landscapes

To quantify the abundance of each contigs, the clean reads were mapped to virome assembly of respective species using Bowtie 2 [51]. Multiple alignment was allowed to obtain maximum sensitivity. The read counts were obtained by FeatureCounts [55] and were normalized and converted to count per million reads (CPM) by R edgeR package [56]. We also used edgeR to conduct differential analysis between different landscape types. Prior to that, each contig, including endogenous and viral, was filtered based on its expression level [57]. Only contigs with sufficiently high expression levels were retained for statistical tests. The contigs with FDR less than 0.05 were considered significantly difference between landscapes. The analysis then focused on viral contigs as defined by the viral identification step.

## Acknowledgements

We are grateful for the help received from Stephanie Gardner, Katherine Odanaka, and Wyatt Shell with collecting bee samples. We thank the Genomic Shared Resource Shared Resource at The Ohio State University Comprehensive Cancer Center, Columbus, OH for generating the virus RNA-seq data reported in this publication. Research reported in this publication was supported by USDA Evans Allen Fund NI231445XXXXG004, NI241445XXXXG004 for HL-B and ZZ, USDA CBG GRANT13202344 for HL-B, USDA award 2020-67014-31557 for MCF, and the NSF Louis Stokes Alliances for Minority Participation (LSAMP) program Ohio Alliance scholarship for JM, the Ohio State University Comprehensive Cancer Center and the National Institutes of Health under grant number P30 CA016058.

## Author contributions

HL-B and SMR designed the experiment. SMR conducted the field studies and provided the samples. MCF and JM prepared samples for RNA-seq. ZZ conducted data analysis and wrote the first draft of the manuscript. All authors reviewed the final manuscript.

